# A wireless centrifuge force microscope (CFM) enables multiplexed single-molecule experiments in a commercial centrifuge

**DOI:** 10.1101/060269

**Authors:** Tony Hoang, Dhruv S. Patel, Ken Halvorsen

## Abstract

The centrifuge force microscope (CFM) was recently introduced as a platform for massively parallel single-molecule manipulation and analysis. Here we developed a low-cost and selfcontained CFM module that works directly within a commercial centrifuge, greatly improving accessibility and ease of use. Our instrument incorporates research grade video microscopy, a power source, a computer, and wireless transmission capability to simultaneously monitor many individually tethered microspheres. We validated the instrument by performing single-molecule force shearing of short DNA duplexes. For a 7 bp duplex, we collected over 1000 statistics of force dependent shearing kinetics from 2 pN to 12 pN with dissociation times in the range of 10-100 seconds. We extended the measurement to a 10 bp duplex, applying a 12 pN force clamp and directly observing single-molecule dissociation over an 85 minute experiment. Our new CFM module facilitates simple and inexpensive experiments that dramatically improve access to singlemolecule analysis.

## Introduction

The power of single-molecule experiments has been well established over the last couple of decades, with various experimental platforms (optical and magnetic tweezers, AFM, and others) enabling unparalleled insights into nanoscale biology and chemistry^1–3^. Unfortunately, single molecule sensitivity often comes at a high price in terms of time, money, and expertise. The vast majority of single-molecule pulling experiments are performed one molecule at a time (with some notable exceptions^4–6^), on sophisticated equipment that often exceeds $100,000. These factors conspire to lower the accessibility of single-molecule techniques to the biomedical researchers that could arguably benefit most from their use.

In previous work, we presented an approach to solve these challenges by using centrifugal force to perform hundreds to thousands of simultaneous single-molecule pulling experiments^7^. In an instrument we call the centrifuge force microscope (CFM), a microscope is rotated such that the optical axis is aligned with the force, enabling observation of many microspheres tethered to a cover glass. Our first prototype was essentially a custom made open-air centrifuge supporting a microscope imaging arm (Figure 1a). While important for establishing proof-of-concept, the prototype had shortcomings including safety, ease of use, and dynamic range. More recently, we developed a new CFM that integrates into a modified benchtop centrifuge (Figure 1b), improving safety, reducing cost, and adding features^8^.

In this work, we introduce a fully wireless and self-contained CFM module that enables use in commercial benchtop centrifuge (Figure 1c). This advance dramatically increases accessibility by eliminating the need for centrifuge modification (and thus a dedicated centrifuge), lowering the cost (below $2,000), and simplifying operation. We have engineered hardware and software necessary to remotely perform video microscopy within a rotating centrifuge. To illustrate application in single-molecule biophysics, we applied shearing forces to short DNA duplexes and observed force and base pair dependent dissociation. It is our hope that this work further democratizes single-molecule experiments, enabling use by a broad range of scientific researchers. Furthermore, with cost effective and straightforward reproduction of our CFM module, the instrument may find uses beyond single-molecule biophysics in areas such as colloidal physics, biological separations, and cell adhesion.

**Figure 1:**
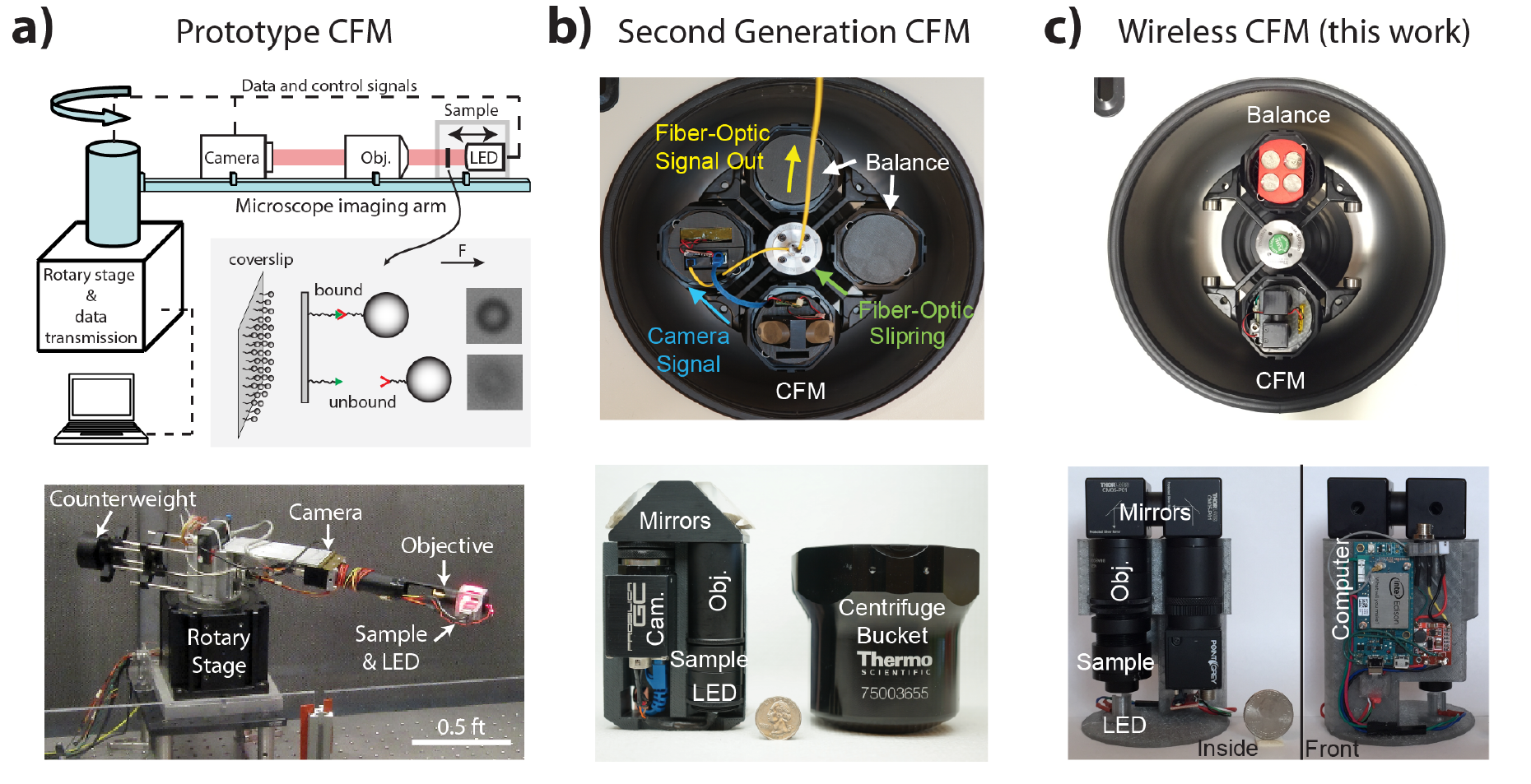
Evolution of the CFM. a) The first prototype used a table mounted rotary stage with a microscope imaging arm and counterweight. b) The second generation instrument integrated into a swinging bucket centrifuge and communicated with an external computer via a connected fiber optic, c) The wireless CFM presented here is completely self contained with onboard power, computing, storage, and wireless data transmission.

## Experimental Section

### Instrument

The CFM module is designed to fit within a commercial swinging bucket centrifuge (Sorvall X1R) with 400mL buckets. To the extent possible, we used commercially available parts to facilitate reproduction by other labs. Labeled images and a complete parts list are given in the Supporting Information (Figure S1 and Table S1). Briefly, the optical assembly is made from aluminum lens tubes (Thorlabs) supporting the camera (Point Grey Flea3), infinity-corrected objective (Olympus Plan Achromat - see Note S1), tube lens (Thorlabs achromatic doublet), turning mirrors (Thorlabs), sample, and light diffuser (Thorlabs). This optical assembly is contained within the 3D printed housing (see File S1 for drawings), which also contains the LED, a lithium polymer battery, single board computer, and supporting circuitry (wiring details in Figure S2).

The module is counterbalanced by quarters arranged in 3 tiers from a 3D printed housing (see File S1 for drawing). The mass of the CFM module is balanced by using the appropriate number of quarters, while the center of mass is balanced by rearranging the positioning of the quarters among 12 possible locations.

### Sample and microsphere preparation

DNA handles were prepared by hybridizing 122 oligos (Integrated DNA Technologies) to a genomic single-stranded DNA, as described in previous work^8,9^. DNA handle D-1 was constructed with a 10 nt 3’ overhang and a 5’ double biotin. DNA handles D-2, D-3, and D-4 contained 3’ overhangs (10 nt, 7 nt, and 6 nt, respectively) with reverse complementarity to DNA handle 1 and also contained a 5’ double biotin. A list of oligos is presented in Table S-2, where the 122 oligos for each DNA handle is comprised of 109 “backbone” oligos, 12 “variable” oligos, and 1 “overhang” oligo.

To prepare the DNA coated beads, we took 40μl of streptavidin microspheres (Thermo Fisher Dynabeads M-270 2.8 μm diameter) and washed twice with 100μl of PBS +0.1% tween 20 using a magnet (Thermo Fisher DynaMag Spin Magnet). Following the wash, the beads were brought to a 10μl volume, and 10μl of the biotin labeled DNA D-1 (~300 pM) was added and reacted for 1 hour on a vortexer to prevent settling of the beads.

### Chamber assembly and preparation

The sample chamber was made by two cover glasses (Warner Instruments) separated by double stick tape (Kaptontape.com) and supported on a flat disc with a central opening. The process of building the chamber is illustrated in Figure S3. The prepared chamber holds 1-10 μL. To prepare the chamber for single molecule experiments, we first flowed in 0.1 mg/mL of Streptavidin (Amresco) in PBS, and allowed to adsorb to the surface for ~1 minute. The chamber was then washed with PBS + 0.1% tween by flowing through four volumes of 50 μL in 30 second intervals. Next, 5 μL of the DNA D-2, D-3, or D-4 (~1.5 nM) was incubated for 5 minutes to allow binding of the biotin labels to the streptavidin surface. Another 4 rounds of 50 μL PBS + 0.1% tween was repeated to remove unreacted DNA. At this point the DNA coated dynabeads were added (typically 5 μL). Finally, the chamber was sealed with high vacuum grease to prevent liquid escape.

### Experimental protocol

The prepared and sealed chamber was securely screwed into the lower lens tube and then screwed into the rest of the optical assembly. With the camera operating from the host computer, we manually adjusted the focus by threading toward the objective and locking into place using a lens tube coupler. Once secure, the 3D housing was closed and the battery and the microUSB plug were connected. At this point, the CFM module was lightly tapped to remove microspheres from the coverglass and then inverted to allow control of the microsphere touch time against the coverglass. Following the touch time, the CFM module was inverted back, and then loaded into the centrifuge bucket. The run was started by a command line script from the host computer (see Note S2 and File S2), and the centrifuge was controlled normally from the front panel to control speed. We typically started image recording before starting the centrifuge.

The centrifuge speed is related to the force on the tethers by F = mω^2^R, where m is the mass of the bead (minus the mass of the medium displaced), ω is the magnitude of the angular velocity, and R is the distance of the microspheres from the axis of rotation. The mass of each microsphere is 6.9 × 10^−12^ g as previously determined^7,8^, and the distance R was measured to be 133 mm. Using these values, we used the following speeds: 2 pN = 445 RPM, 6 pN = 770 RPM, 8 pN = 890 RPM, 10 pN = 995 RPM, and 12 pN = 1090 RPM.

### Data analysis

Images were analyzed by first identifying tethered beads in the first movie frame following acceleration. Tethered microspheres were visually selected based on focus (see Figure 3a). Double tethers (though rare) were eliminated as they were observed, typically seen as a lateral shift when the first tether breaks. Tethered microspheres were tracked throughout subsequent frames and the number of microspheres being released at each frame was recorded.

### Safety considerations

These experiments involve atypical use of a centrifuge, which can involve enormous kinetic energy. While commercial centrifuges are generally designed to contain any catastrophic failure, great care must be taken to ensure that: 1) the CFM module is properly balanced, 2) that the masses or speeds used do not exceed those recommended by the manufacturer, 3) that all components are secured within the buckets.

## Results and Discussion

We developed a wireless CFM module to improve the accessibility and reduce the cost of multiplexed single molecule experiments. The module was designed to be fully compatible with existing commercial centrifuges, and we used the Sorvall X1R swinging bucket centrifuge as our model system (Figure 2a). Following a similar optical design as previous^8^, we additionally implemented an onboard computing and data transmission strategy to allow the CFM to fit within a single bucket, and to enable use without modification of the centrifuge (Figure 2b and 2c). We chose to use the Intel Edison as our onboard computer due to its small size, low cost, and integrated WiFi capabilities. We interfaced the onboard computer with a small USB machine vision camera (Point Grey Flea3) and developed a 3D printed housing to secure the various optical and electrical components (see File S1). The resulting CFM module can be built for ~$1,000-$2,000 depending on specific camera and objective choice (see Table S1 and Note S1). All parts are commercially available, with the exception of one custom machined thread adapter for which we have offered alternatives (see Note S1), and the 3D printed housing. However, we have made available the drawing files for the 3D printing, and capable 3D printers are now widely available, even at many public libraries.

We additionally developed software and procedures to facilitate the fabrication and use of the CFM (See Supplemental Information). Briefly, the onboard computer was loaded with a Linux based operating system, modified to enable USB transmission of large files, and loaded with the Point Grey camera drivers. Control of the onboard computer was then performed remotely from a host computer (See Note S2). The communication scheme for normal operation is outlined in Figure 2d, requiring only 2 steps from the user: 1) connecting the CFM with its battery power source, 2) running a program from the host computer command line dictating the number of images and frame rate. These instructions were wirelessly received by the CFM and executed. The centrifuge was controlled through the front panel to control speed, acceleration, temperature, and time. At the end of the run, the image files were wirelessly transmitted back to the host machine and erased from the onboard memory.

Using this setup, we recorded images of 2080 × 1552 pixels to accommodate >2000 microspheres (2.8μm diameter) with a 40x objective and >8000 for a 20x objective (Figure 2e). We used a pixel depth of 8 bits, yielding an image size ~3.2 MB with the first 4 pixels encoding the camera generated timestamp. The onboard computer contains 4GB of built in (eMMC) storage, which leaves 1-2 GB, or about 300-600 frames, of free memory for file storage. We found that saving images to RAM and later saving them to the eMMC resulted in faster maximum frame rates (up to 10 fps at full resolution without skipped frames) due to less computational overhead, but reduced the total number of frames to ~250 due to the lower storage capacity (1 GB). For frame rates slower than 1 fps, we grabbed 1 image at a time with a pause in software, achieving run times approaching 1.5 hours on a single battery. The post experiment transfer is largely dictated by WiFi speeds, and in our hands we found a transfer speed ~5 MB/s (1.4 fps on average).

**Figure 2:**
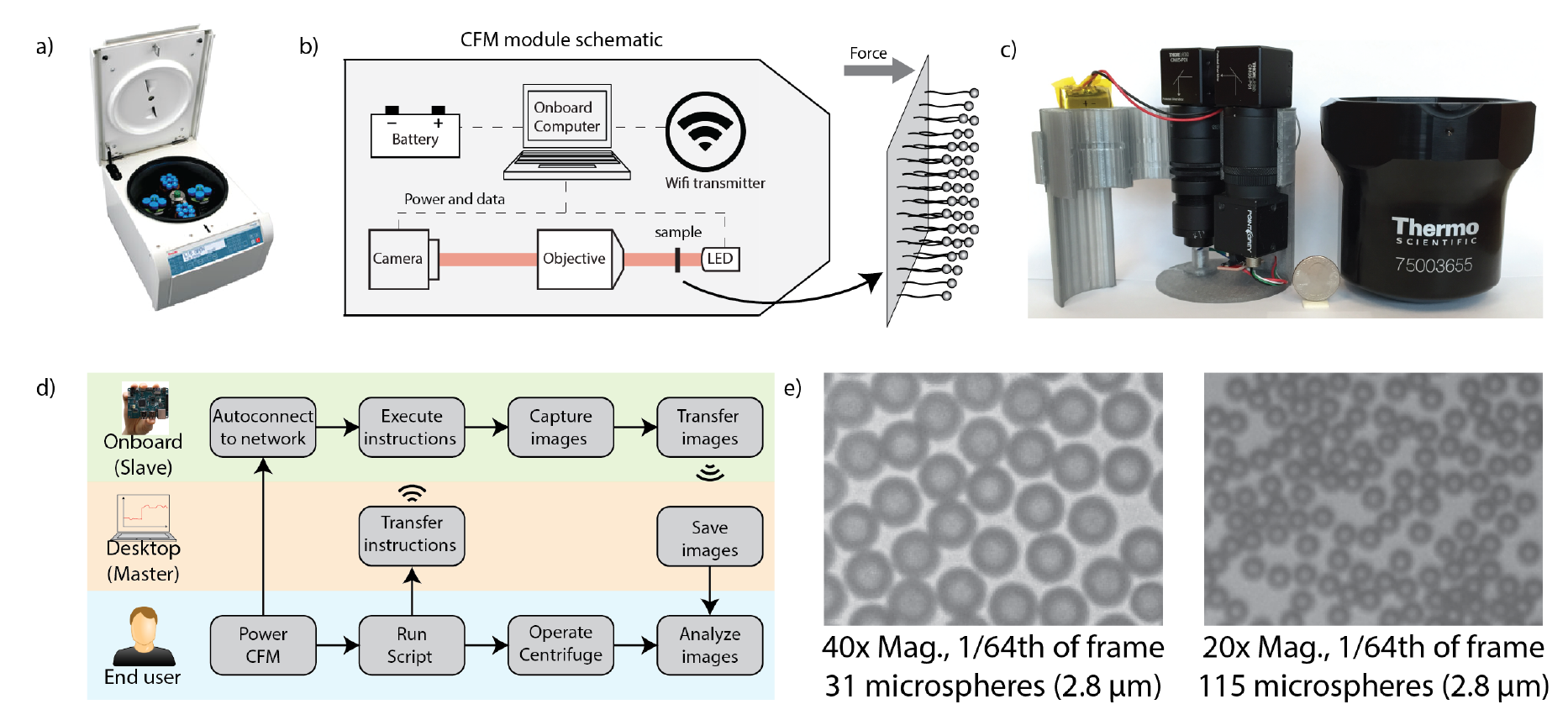
Instrument design and validation. a) Our model benchtop centrifuge (Sorvall X1 series), b and c) The CFM module is designed to fit in a swinging bucket, and contains a miniature microscope, a battery, and a computer with wifi capabilities, d) Flow chart for CFM control software, e) 260×194 pixel images from the CFM module at 40x and 20x magnification, representing 1/64^th^ of the full frame size.

To demonstrate single-molecule manipulation on the wireless CFM module, we performed experiments to shear short DNA duplexes. We constructed double biotin labeled DNA handles with short complementary overhangs. The two DNA handles were separately reacted with streptavidin beads and a streptavidin coated coverglass to enable single molecule tethers connected by a short duplex (Figure 3a). We made beads with various dilutions of the DNA until we found conditions that consistently produced <15% attachment frequency, which is desired to minimize multiple tethers. Using this arrangement we incubated DNA loaded beads against the surface for ~1 minute to enable formation of the duplex, and then started the centrifuge to attain a constant desired velocity.

We first performed constant force pulling experiments on a 7bp duplex. As expected, we found the distribution of rupture times to follow an exponential decay (Figure 3b). With the large number of statistics, we were additionally able to independently analyze multiple runs under the same conditions to assess repeatability (Figure 3b inset). From fitting five individual distributions, we found a mean dissociation time constant for 12 pN of 14.8 seconds with a standard deviation of 1.8 seconds. As a comparison, fitting the cumulative distribution of the combined data gave a time constant of 14.9 ± 0.1 seconds.

We expanded the data by repeating the measurement at different force levels over the range from 2 pN to 12 pN, with time constants increasing for smaller forces (Figure 3c).We fit these results with the Bell-Evans model for force activated dissociation^10,11^ where the force dependent off rate is described as k_off_(f) = k_off_ exp(f/f_β_), where k_off_ is the thermal off-rate, f is the applied force, and f_β_ is the characteristic force scale of the interaction. f_β_ can equivalently be expressed as k_B_T/x, where k_B_ is the Boltzmann constant, T is the absolute temperature, and x is the length scale describing the separation of the bound and unbound states. Using this model, we found a thermal off-rate k_off_ of 0.004 ± 0.002 s^-1^ and a force scale f_β_ of 4.5 ± 1.0 pN.

**Figure 3:**
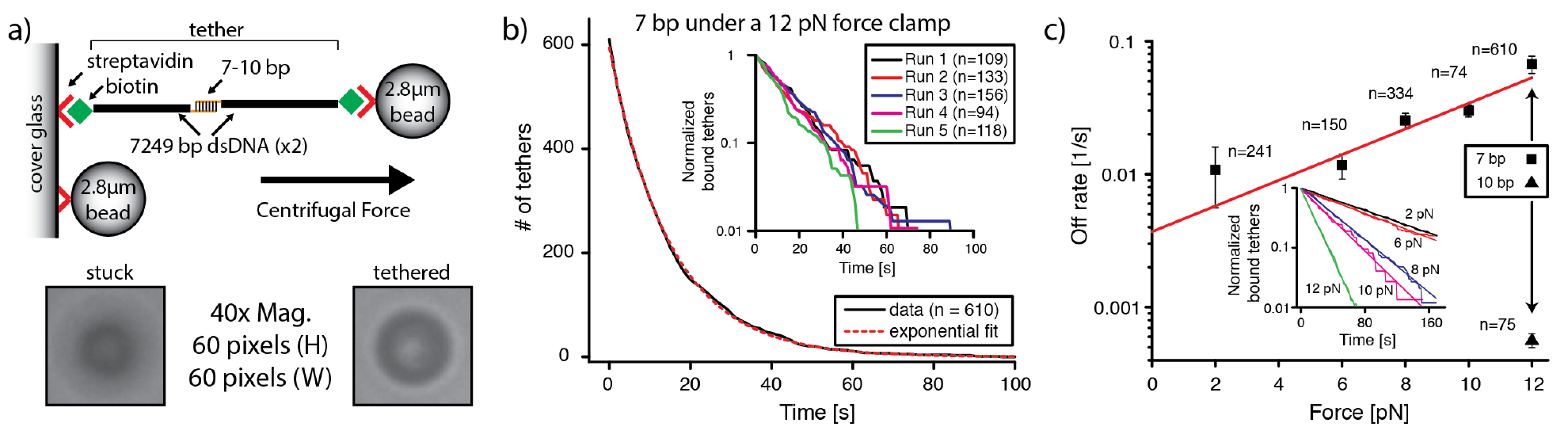
Force shearing of short DNA duplexes. a) The experimental setup consists of microspheres tethered to a cover glass by biotin labeled DNA handles, with a short central duplex dissociating under applied centrifugal force. b) For the 7bp duplex under a 12 pN force clamp, we collected over 600 statistics and found a dissociation time constant of 14.9 seconds. By independently analyzing the five independent runs comprising the data, we found good repeatability and estimated an uncertainty of ~15% in the time constant (inset). c) Force dependent dissociation of the 7 bp duplex (squares) showed increasing off-rate as a function of force, with a fit to the data (line) providing a solution off rate of 0.004 s^−1^ and a force scale of 4.5 pN. A comparison with a 10 bp duplex shows a dramatically slower off-rate (triangle). Individual data for each force is shown in the inset. Error bars were determined based on the standard deviation of multiple independent experiments.

We additionally investigated a 6 bp duplex and a 10 bp duplex at a constant force of 12pN. As expected, we observed markedly slower dissociation for the 10 bp duplex, observing dissociation over an 85 minute experiment to determine a time constant of 28 ± 1 minutes. In attempting to measure the 6 bp duplex, we did not observe significant tethers over several runs, presumably because the dissociation time scale decreased too dramatically. Over this relatively narrow range of lengths, the force scale does not change dramatically^12^, so that our 10 bp off rate can reasonably be extrapolated to estimate thermal off-rate. Based on this, we estimate ~ 4 × 10^−5^ s^−1^, which corresponds to a dissociation time constant of ~ 7 hours.

In comparing our results with others, we note that most if not all DNA shearing experiments have used duplexes 10 bp or larger^12-14^, and one has even predicted that 8 bp or less will not be measureable^15^. The sequence of our 10bp duplex was chosen to match previous work^12^, while the shorter duplexes were truncated from that sequence. In comparison to that work, we found a force scale for the 7bp duplex that agrees within error, but our off-rates are notably slower than those predicted in that work. For the 7 bp and 10 bp duplexes, they predict off rates of 0.3 s^−1^ and 0.01 s^−1^, respectively, which are approximately two orders of magnitude faster than our measurements.

There are a few factors that likely contribute to this difference. Our 10 bp duplex, while of identical sequence, has flanking bases on both sides that contribute base stacking interactions. Calculations using Mfold^16^ estimate that these stacking interactions contribute an additional 1.5 to 2.7 kcal/mol to the stability of the 10 bp duplex, suggesting an off-rate that should be one to two orders of magnitude slower. Even using the conservative (1.5 kcal/mol) adjustment would put our measurements within error of theirs^12^. Similar corrections would apply for our shorter duplexes as well. Another (less significant) contribution to the observed difference may be due to an overestimation of the off-rate in ref [12]. They approximated that the force-loading rate was dictated by the tether elasticity at rupture, which overestimates the loading rate (and thus the off-rate) since much of the experimental time is spent reaching that loading rate.

## Conclusions

As we have shown here, the wireless CFM module is capable of multiplexed singlemolecule experiments with throughput that compares favorably with some of the highest throughput methods^4,5^. Our modular design allows unprecedented accessibility to singlemolecule experiments by leveraging common benchtop centrifuges with buckets of 400 mL or larger. The cost of our module is one to two orders of magnitude less expensive than many competing single-molecule technologies, and up to 100,000 times cheaper if one considers cost per pull compared to optical tweezers, for example. The simplicity of operating our CFM module makes it amenable to use by a wide range of researchers including undergraduates (such as one author of this paper). Here we have provided all of the necessary designs, instructions, and software to make reproduction of this system straightforward.

The centrifuge force microscope is still an instrument in its infancy, but has already undergone significant improvement here and in previous work. Still, further development work is needed to improve some of its current limitations. Integration of motorized components to control stage positioning in 3-axes would benefit sample setup, focusing, and tracking. Flow control would enable changing buffers and reagents during experiments for more complex experiments. Further improvements in batteries, single chip computers, and WiFi protocols will enable longer experiments, faster data transmission, and real-time data analysis.

While the CFM is unlikely to supplant existing single-molecule technologies, we believe it occupies an important niche for multiplexed single molecule experiments where perhaps low cost and ease of use are more important than sub-nanometer and submillisecond resolutions. With this work, it is our hope that this CFM module will encourage more widespread single-molecule experimentation by biomedical researchers, and lead to new and exciting avenues of research.

## Acknowledgements

The authors thank Alan Chen and Pan T.X. Li for their financial support and mentorship role for T.H. We thank Eric Warnke for vital software, hardware, and programming support, Arun Chandrasekaran for preparing DNA handles, and Andreas Karl and Thermo Fisher Scientific for providing materials and support for the centrifuge. We acknowledge the SUNY STEM Research Passport program for financial support of D.P., and Combined Funding Application (University at Albany-RNA Institute Capital Project #X765) from the State of New York. We thank Wesley Wong, Darren Yang, Maria Basanta Sanchez, and Sri Ranganathan for useful discussions.

## Author Contributions

TH designed, built, and programmed the instrument and performed experiments. DSP developed protocols and performed experiments. KH conceived the project, analyzed the data, and wrote the paper.

